# A Block-Capable and Module-Extendable 4-Channel Stimulator for Acute Neurophysiology

**DOI:** 10.1101/2020.01.28.923474

**Authors:** Adrien Rapeaux, Timothy G. Constandinou

**Affiliations:** Centre for Bio-Inspired Technology, Imperial College London, SW7 2AZ, UK; Department of Electrical and Electronic Engineering, Imperial College London, SW7 2BT, UK

## Abstract

**Objective:** This paper describes the design, testing and use of a novel multichannel block-capable stimulator for acute neurophysiology experiments to study highly selective neural interfacing techniques. This paper demonstrates the stimulator’s ability to excite and inhibit nerve activity in the rat sciatic nerve model concurrently using monophasic and biphasic nerve stimulation as well as high-frequency alternating current (HFAC).

**Approach:** The proposed stimulator uses a Howland Current Pump circuit as the main analogue stimulator element. 4 current output channels with a common return path were implemented on printed circuit board using Commercial Off-The-Shelf components. Programmable operation is carried out by an ARM Cortex-M4 Microcontroller on the Freescale freedom development platform (K64F).

**Main Results:** This stimulator design achieves *±*10 mA of output current with *±*15 V of compliance and less than 6 µA of resolution using a quad-channel 12-bit external DAC, for four independently driven channels. This allows the stimulator to carry out both excitatory and inhibitory (HFAC block) stimulation. DC Output impedance is above 1 MΩ. Overall cost is less than USD 450 or GBP 350 and device size is approximately 9 cm *×* 6 cm *×* 5 cm.

**Significance:** Experimental neurophysiology often requires significant investment in bulky equipment for specific stimulation requirements, especially when using HFAC block. Different stimulators have limited means of communicating with each other, making protocols more complicated. This device provides an effective solution for multi-channel stimulation and block of nerves, enabling studies on selective neural interfacing in acute scenarios with an affordable, portable and space-saving design for the laboratory. The stimulator can be further upgraded with additional modules to extend functionality while maintaining straightforward programming and integration of functions with one controller.

## 1. Introduction

Acute *Ex-Vivo* and *In-Vivo* Experiments are a cornerstone of research in neurophysiology as they enable a quick exploratory experimental cycle. This reduces the risk of failure when committing to lengthy chronic *in-vivo* experiments before an animal model is well-understood by the experimenter.

A large number of stimulators are available from commercial and academic sources, however these are generally designed for specific applications. Work in academia has often focused on implantable systems for awake-behaving animal neurophysiology [1, 2, 3] or interfacing with multi-electrode arrays (MEAs) [4, 5], with few designs dedicated to acute *in-vivo* or *ex-vivo* neurophysiology [6, 7, 8].

In addition to conventional excitatory stimulation, there has been increasing interest in the use of HFAC block for inhibitory stimulation, resulting in a powerful and versatile toolset for neuromodulation. Notable applications are in the emerging Electroceuticals (also referred to as Bioelectronc Medicine) [9, 10, 11, 12] and Functional Electrical Stimulation [13, 14, 15] fields. Work on HFAC block has been exploratory in nature, with experimental setups used to modulate waveforms or combine multiple current sources in order to achieve specific effects [16, 17, 18, 19, 20].

HFAC block requirements are orders of magnitude above those of conventional stimulation, with large output currents to reach block thresholds of up to 10 mA at high bandwidths of 50 kHz or more, though lower bandwidth 5-10 kHz waveforms are used more often [21, 15, 14, 12]. This requires high compliance voltages of *±*10 V or more as even cuff electrodes with large contacts have impedances in the kilo-ohm range [22]. In addition to this, precise control of stimulation timing and DC leakage is required in order to prevent any damage to nerves resulting from bringing the polarization voltage of electrodes outside of the water window [23, 24, 25] during application of block. For stimulators that provide enough compliance voltage [7, 2, 26, 27], other limitations can include low output DAC resolution or full-scale output current which will limit the ability of the user to carry out standard nerve function tests such as stimulation strength-duration curves, or reach block thresholds in typical targets such as the rat sciatic or vagus nerves. Many stimulators designed as single chips for implantable systems are further limited by maximum pulse width [26, 28] which can limit the frequencies of block used or the method used to deliver conventional stimulation, which may rely on longer pulse widths [29]. An older design such as the Neurodyne [30] could be an excellent and affordable HFAC block stimulator due to its high output current and compliance but is limited by the dynamics of its transformer core when long low-amplitude pulses are required for conventional stimulation. As such stimulators designed for exploratory neurophysiology need to be versatile enough to provide both high current output, high voltage compliance and enable complete control of the waveform for both excitatory and inhibitory stimulation, allowing researchers to combine multiple stimulation methods such as the ones described in [31] and [32].

Commercially available stimulators such as the DS3 (Digitimer) or PlexStim (Plexon) do not generally combine high compliance, high output current, and multiple output channels without significant financial and space investment in the lab. Stimulus Isolators such as the DS5 (Digitimer), A360 (World Precision Instruments) or Model 2200 (A-M Systems) which are V-to-I converters have been successfully used for block but are generally single-channel, regularly require calibration to prevent DC leakage due to their open-loop nature, and may have limited bandwidth at HFAC block frequencies [33]. They may also present an integration and timing challenge for users who require accurate timing between different stimulation channels, are expensive and require auxiliary hardware in the form of external Arbitrary Function Generators. To the best of the authors’ knowledge, no multi-channel stimulators have been designed in academia or industry with high-frequency block as an application. The resulting bulkiness and cost of equipment can be prohibitive in space or budget-constrained laboratories, especially when taking into account the additional investment needed for life support equipment: breathing and anaesthesia apparatus for *in-vivo* [34], saline buffer processing and maintenance setups for *ex-vivo* [12, 35]), and any recording equipment, filters and dataloggers [6].

Based on these observations the following list of features and constraints were used to guide the design of a stimulator meeting all requirements for acute excitatory and inhibitory stimulation:

- High Compliance Voltage to accommodate kilo-ohm scale load impedances
- High Full-Scale Current to reach potentially high nerve fibre block thresholds
- High Current resolution to enable users to carry out precise stimulation or output arbitrary current waveforms without distortion
- High Bandwidth to provide high-frequency block waveforms without distortion
- Multiple independent channels to stimulate multiple areas of a nerve concurrently
- High stimulation timing accuracy, versatility and reliability
- Automatic current output channel self-calibration
- Real-time electrode voltage monitoring to enable the system to correct for charge imbalance over long-term delivery of block
- Small size corresponding to development board dimensions
- Versatile interfacing options to external computers or TTL-triggered equipment
- Battery Operation with at least 8 hour lifetime on one charge

In addition to these targets, special consideration was given to flexible programming and scripting to give the user complete control over the operation of the stimulator. Additional feature extension was made possible by using the Freescale K64F microcontroller development platform, notably for the addition of a neural recording module on top of the stimulation module. This would enable complete integration between stimulation and recording, allowing closed-loop control of stimulation using external computers for processing and control, and straightforward embedding of stimulation parameters for each stimulation event in recording metadata. A photograph of the assembly with the stimulator module on top of the microcontroller development board is shown on Figure 1.

**Figure 1:**
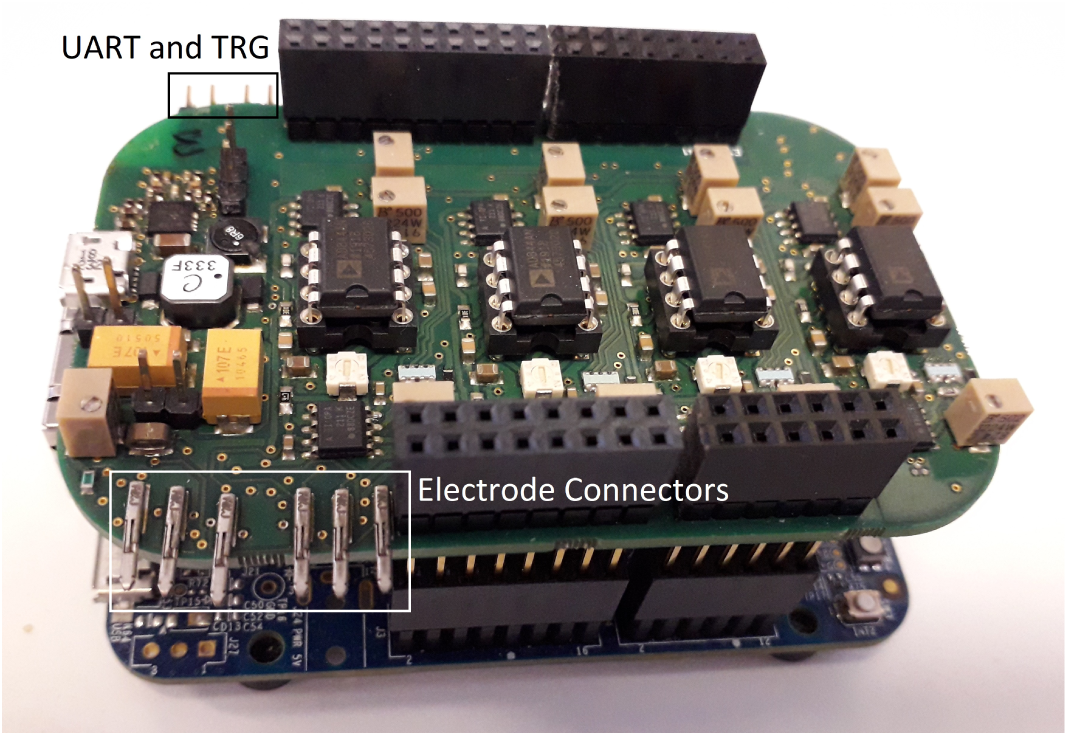
Stimulator board on top of the Freescale K64F Microcontroller development board, with labelled input-output pins.

## 2. Principle of Operation

### 2.1. Module Power and Data Isolation

The stimulator system was designed to ensure that in experimental setups there is at most a single earth point in contact with the nerve tissue. As neural signal acquisition devices often ground the experimental setup with earth ground to reduce noise, the stimulator must be battery powered (see Figure 9). Separating the microcontroller and stimulator power domains allows recording neural signals using devices on the microcontroller power domain, as shown Figure 2. This enables isolated recording and stimulation using a single device. The PC used to control the stimulator is electrically isolated from the stimulator itself, as other devices may also be connected to the same computer for signal acquisition or experimental automation, or the computer may itself be connected to earth ground in the case of a desktop. Such a setup prevents ground loops, stimulation interfering with recording and improves setup flexibility.

**Figure 2:**
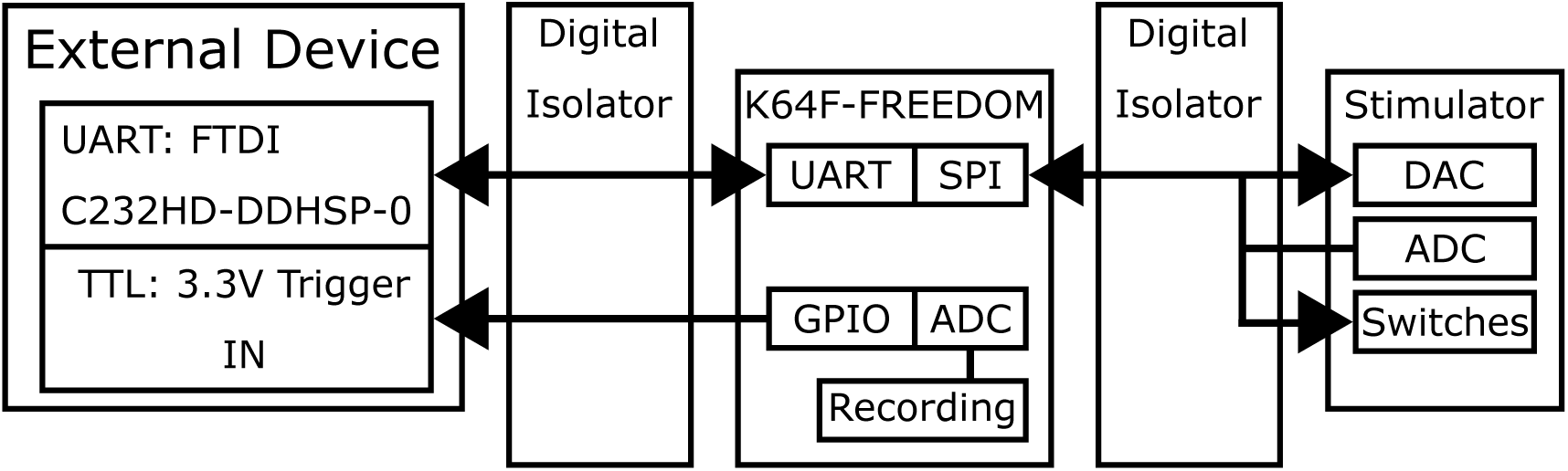
Power domain isolation and inter-device communication interfaces.

The power supply was chosen for maximum flexibility. Using 5 V as an input voltage for the system allows laptops and portable phone and tablet power banks to be used to supply power, providing long operating life which can be easily sourced. To achieve *±*15 V compliance at the stimulator output, the output voltage of the power supply was chosen as *±*18 V which allows a wide range of operational amplifiers to be used. The ADP5070 dual output switch-mode power supply provided a compact and efficient solution for this with manageable output ripple. Manufacturer recommended values were used for all power supply components and assembled using the recommended layout. High frequency noise at the output was filtered using ferrite beads, and shielded inductors were used to reduce EMI emissions that could interfere with sensitive nodes in the Howland Current Pump. For precision voltage references such as ADC references, Low drop-out (LDO) components were used to filter out noise from higher voltage supplies. To provide an isolated power supply to the microcontroller power domain, a NXJ2S0505MC-R7 (MURATA Power Solutions) isolated 5 V to 5 V DC-DC converter was used.

To allow devices on different power domains to transfer data, the microcontroller’s Serial Peripheral Interface was used together with a 6-channel digital isolator as shown Figure 2. Communication to the PC is carried out by passing UART peripheral signals through a similar 4-channel digital isolator. Noise emissions from communication were kept to a minimum by segregating digital interface components close to one edge of the PCB while the opposite edge was used for analog input-output.

### 2.2. Current Source

The Howland Current Pump (see Figure 3) was chosen as the stimulator’s current source circuit due to it not requiring an H-bridge to provide bidirectional current sourcing and sinking, high speed and a lower number of control signals that must go through a digital isolator to drive the circuit, as only the DAC must be digitally driven during stimulation. The output current equation is as follows for this circuit:

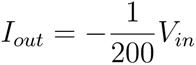

**Figure 3:**
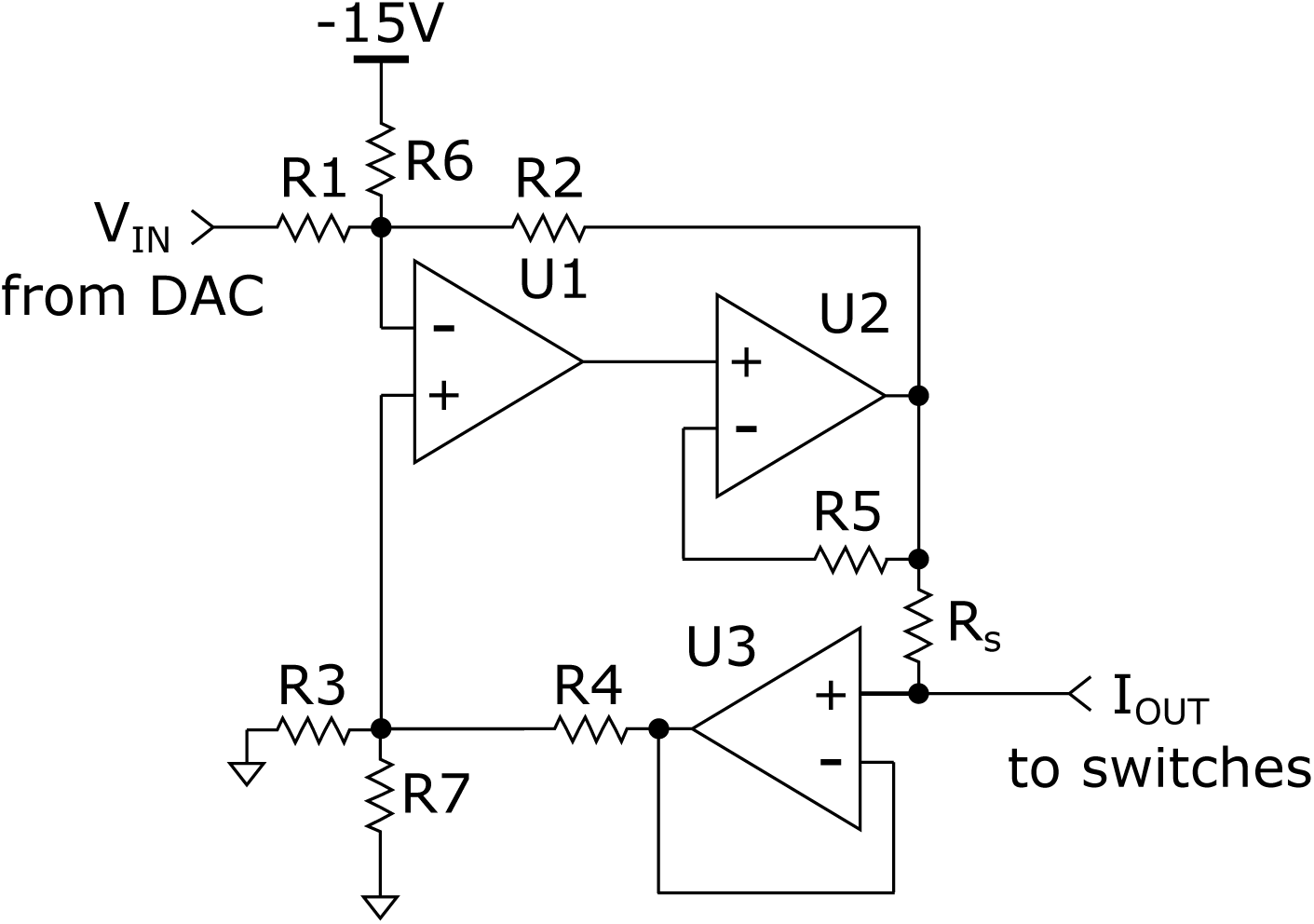
Circuit schematic for the Howland current pump used as the current source for the stimulator channels.

A compound amplifier solution in U1 and U2 was chosen to benefit from the low offset of U1 and the output capability of U2, which is a current feedback amplifier. U2 can be placed in either a non-inverting or inverting configuration. The inverting configuration is slightly faster at the cost of significantly higher power consumption for U1 which would source current into the feedback resistors of U2. For this reason a non-inverting configuration was chosen. As the feedback signal depends on the current flowing into R*_S_*, any leakage beyond this point will not be compensated for and therefore U3 is used as a buffer to prevent current leakage. As the half-scale output of the DAC is 2.048 V, an offset adjustment is implemented with R6 such that when the DAC output is at half-scale the voltage on the inverting input of U1 is 0 V, leading to 0 A current output. R1, R2, R3 and R4 are closely matched resistors packaged in a single device (Vishay ACAS, 0.05% relative tolerance), and additional 50 Ω potentiometers (3224 series, Bourns, not shown) are used in series for precision matching. R6 is matched to R7 using a 5 kΩ potentiometer. Target values for components are listed in Table 1.

**Table 1:** Component values for the current output channel.

| Component | Value | Component | Value |
| --- | --- | --- | --- |
| R1 | 10 kΩ | R6 | 64.2 kΩ |
| R2 | 1 kΩ | R7 | 64.2 kΩ |
| R3 | 10 kΩ | U1 | OPA211 |
| R4 | 1 kΩ | U2 | AD844 |
| R5 | 1 kΩ | U3 | OPA211 |
| DAC | MAX5580AEUP+ | R<sub>S</sub> | 20Ω |

### 2.3. Self Diagnostics

The ability of a neural stimulator to detect out-of-compliance conditions, correct for drift over time and temperature, and measure residual voltage across stimulation electrodes is becoming standard in commercial hardware and these features were included in the form of an acquisition chain with an instrumentation amplifier and an external ADC on the stimulator power domain, combined with a system of switches to connect current output channels and the monitoring system to electrodes, shown Figure 4 and with component values detailed in Table 2. As it can be useful to measure stimulation electrode voltage with respect to an electrochemical reference in the nerve bath during stimulation, the dedicated REF connector is available for this. This can be used to determine when stimulation electrodes are polarized outside of the ‘water window’ for example, indicating that water splitting reactions can occur and change local pH, potentially affecting the nerve tissue. A dummy electrode implemented as a resistor and capacitor in series was also included with the ability to short the capacitor through a switch, for calibration purposes.

**Figure 4:**
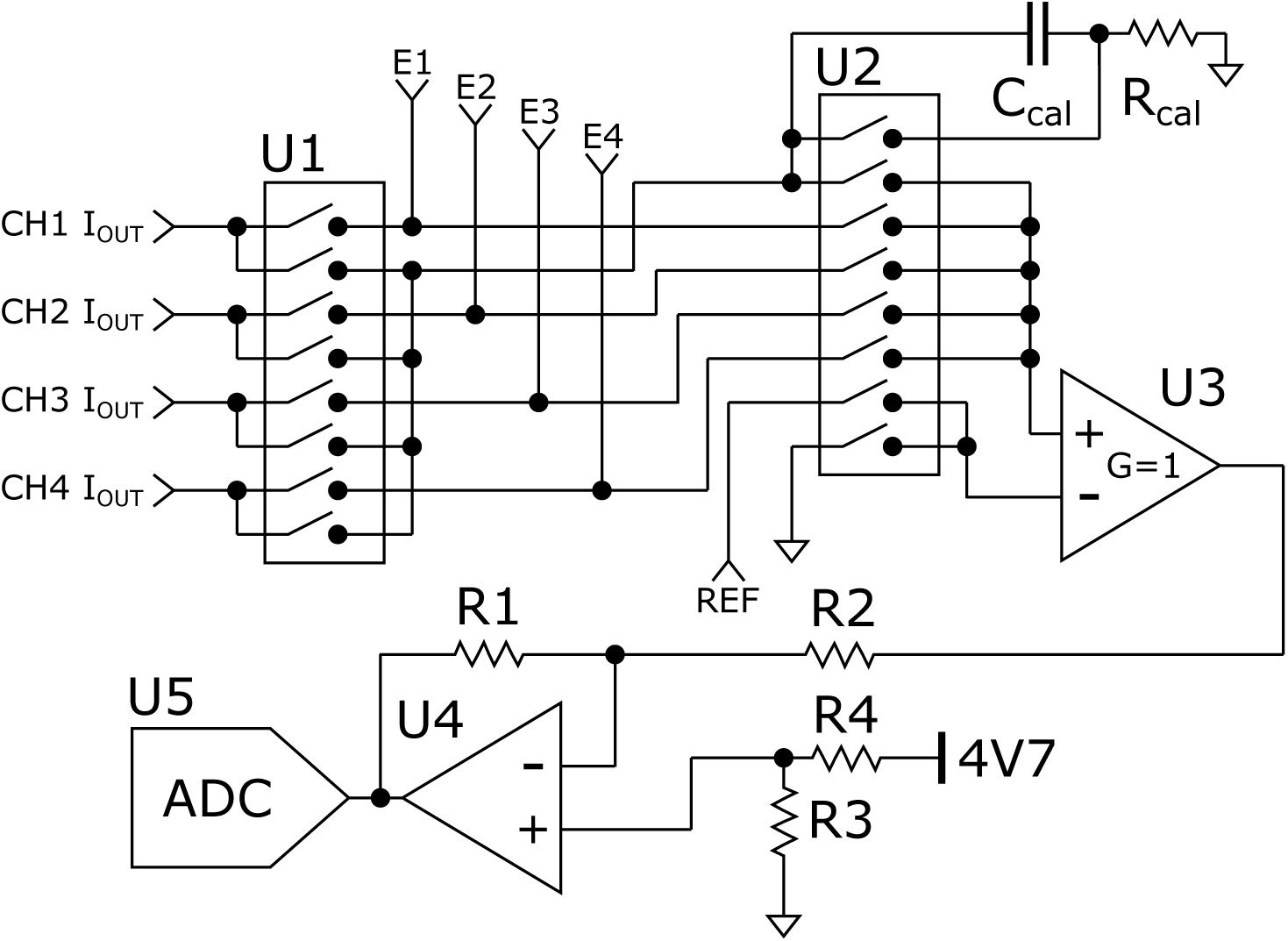
Circuit schematic for the routing and diagnostic module of the stimulator. CH1-CH4 refer to the output nodes for each stimulation channel. E1-E4 refer to individual electrode connectors for each stimulation channel. REF refers to a reference electrode connector for when the monitoring circuit should measure voltage between an electrochemical reference such as silver-silver chloride and a stimulation electrode, rather than using system ground as a reference.

**Table 2:** Component values for the diagnostics module.

| Component | Value | Component | Value |
| --- | --- | --- | --- |
| R1 | 1 kΩ | U1 | ADGS5414 |
| R2 | 9.09 kΩ | U2 | ADGS5414 |
| R3 | 2 kΩ | U3 | AD8422 |
| R4 | 3.24 kΩ | U4 | OPA211 |
| R<sub>cal</sub> | 1 kΩ | U5 | LTC2313ITS8-14 |
| C<sub>cal</sub> | 100 nF |  |  |

### 2.4. Address-Event Representation of Stimulation

Precisely-timed stimulation is required when multiple sources of stimulation are active simultaneously, for example when current steering or combining stimulation events in multiple locations on the nerve. In order to guarantee precise timing and avoid using processor interrupts which can delay operation for other tasks such as responding to commands or processing recording input during stimulation, the SPI link to the stimulator’s quad-channel DAC was driven using microcontroller peripherals only. By triggering the K64F enhanced Direct Memory Access (eDMA) peripheral using the Periodic Interrupt Timer (PIT) peripheral, it is possible to output complex waveforms through the external SPI-controlled DAC of the stimulator. eDMA channels can be used to both send new output values to the external DAC and change the value of the PIT Load Register, changing the cycle time of the triggering PIT timer as shown in Figure 5. The Kinetis K64 microcontroller has 4 PIT channels and 16 eDMA channels and can therefore drive 4 stimulator channels independently and with no processor involvement during stimulation. When it is required to deliver more stimulation events than it is possible to store in memory, the eDMA peripheral’s scatter-gather feature can be used to have certain eDMA channels track and change other channels’ behaviour as a stimulation protocol progresses. This is useful especially when delivering neural block waveforms which due to their high-frequency nature require many stimulation cycles to be output quickly and with precise timing. Long pulses can also be output thanks to the PIT timer’s 32-bit wide timer register. In this design the PIT is clocked at its maximum frequency of 60 MHz, which translates to a maximum timer period of approximately 71 seconds, and the same maximum stimulation phase duration.

**Figure 5:**
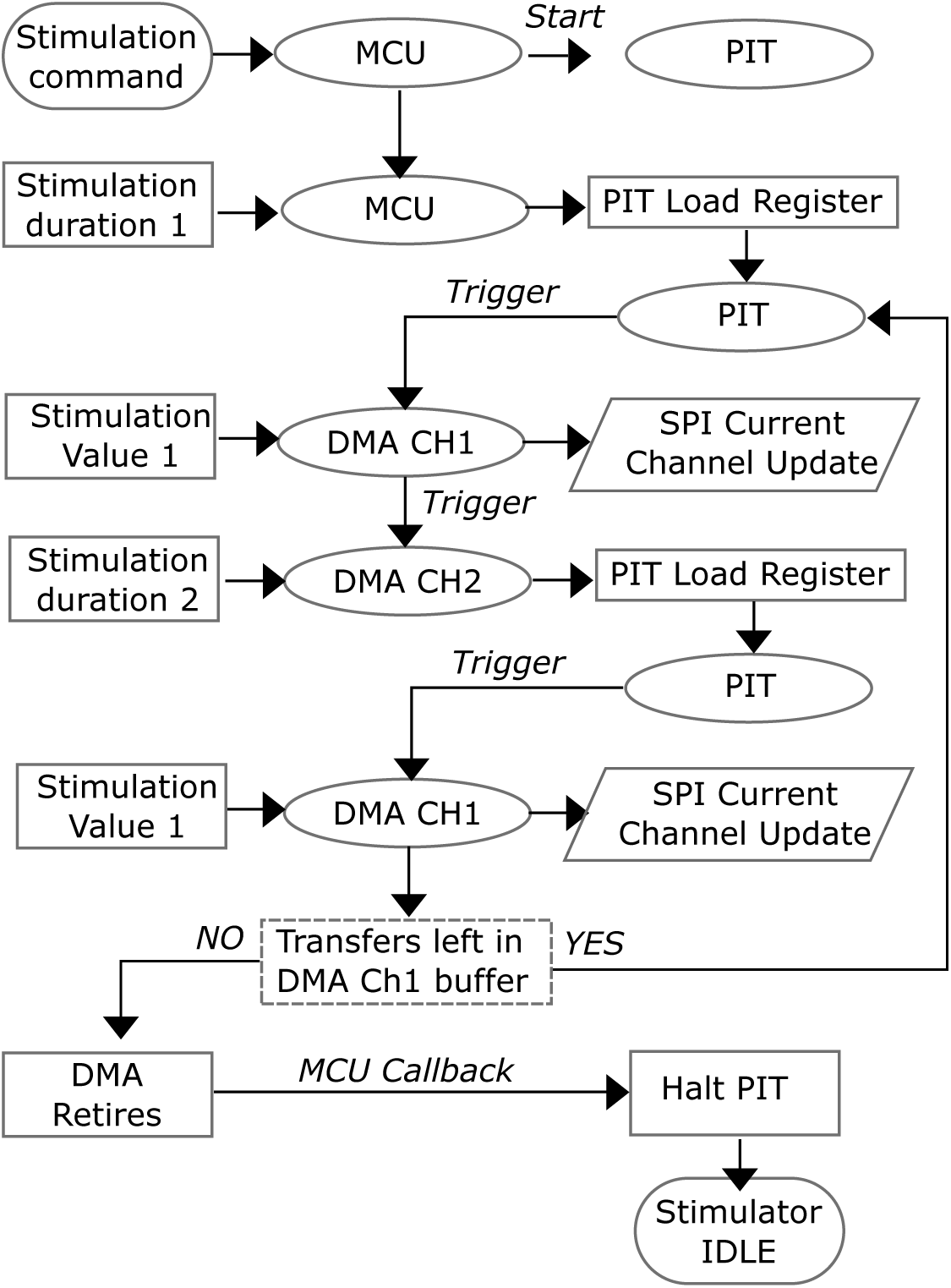
Flowchart describing how stimulation is carried out without processor intervention. The processor is only used at the start and end to configure the eDMA and timers and to halt the timers and reset the eDMA when timing is not critical.

## 3. Electrical Performance Results

### 3.1. Output Impedance over output voltage and frequency

To characterise the stimulator’s ability to regulate current output at different output voltages and output signal frequencies, two tests were carried out. To measure compliance, output node voltage was swept from −18 V to +18 V in 100 mV steps using a Keithley 2635B Sourcemeter at 10 mA current output from the stimulator both for sinking and sourcing cases. Keithley received current was measured as a function of stimulator output node voltage to determine the compliance limits of the stimulator, with results on Figure 7 showing voltage compliance of *±*15 V before noticeable output error.

**Figure 7:**
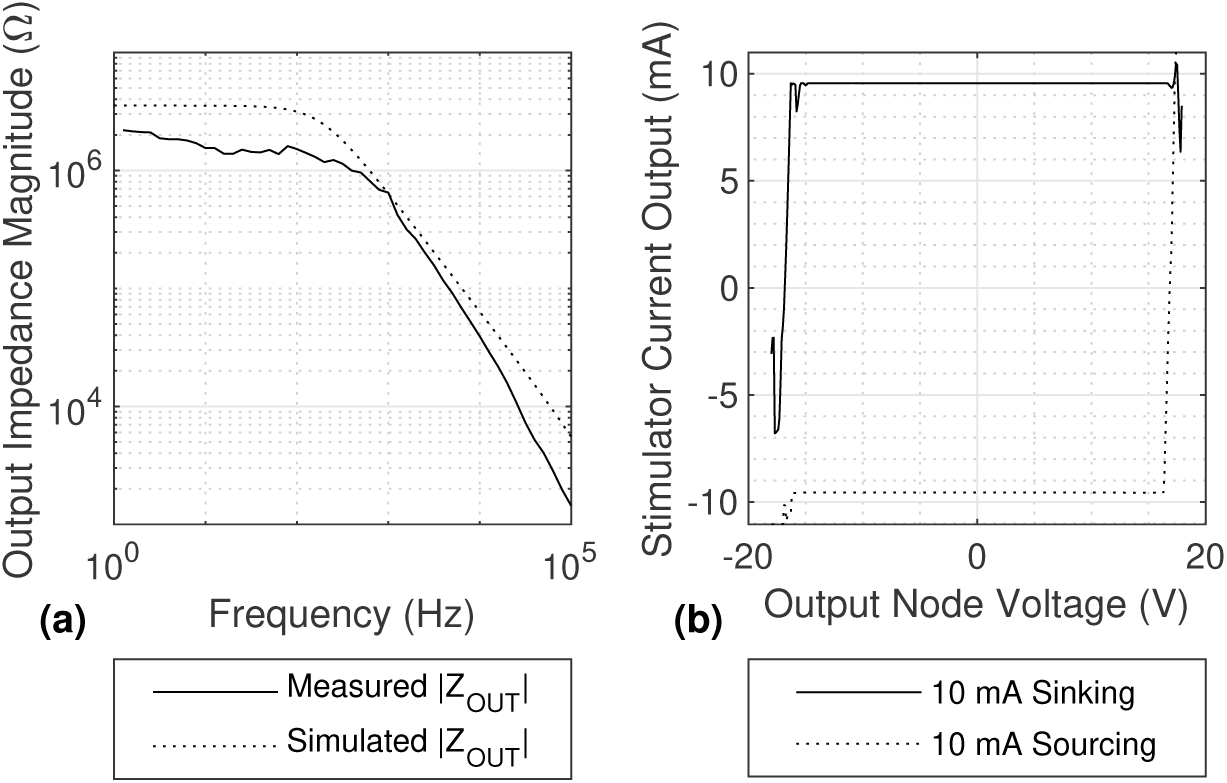
(a) Output impedance magnitude over frequency for one current output channel of the stimulator, comparing simulated and measured results; (b) measured current output of the stimulator when sweeping the output node voltage in maximum current sourcing and sinking scenarios.

To measure output impedance magnitude over frequency, the setup shown on Figure 6 was used. The output impedance plot is shown Figure 7 with a cut-off frequency of approximately 1 kHz. The DC output impedance magnitude is approximately 2 MΩ. Assuming the series resistance of the arbitrary waveform generator (AFG3102, Tektronix) is negligible compared to the 15 kΩ measurement resistance, the latter and the output impedance can be considered in parallel. The output impedance magnitude was calculated using the following equation:

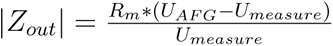

where *U_AFG_* is the amplitude of the AFG output sine wave and *U_measure_* is the amplitude of the sine wave measured at the terminals of the measuring resistance. A simulation of the circuit was also carried out in LTspice XVII for comparison.

**Figure 6:**
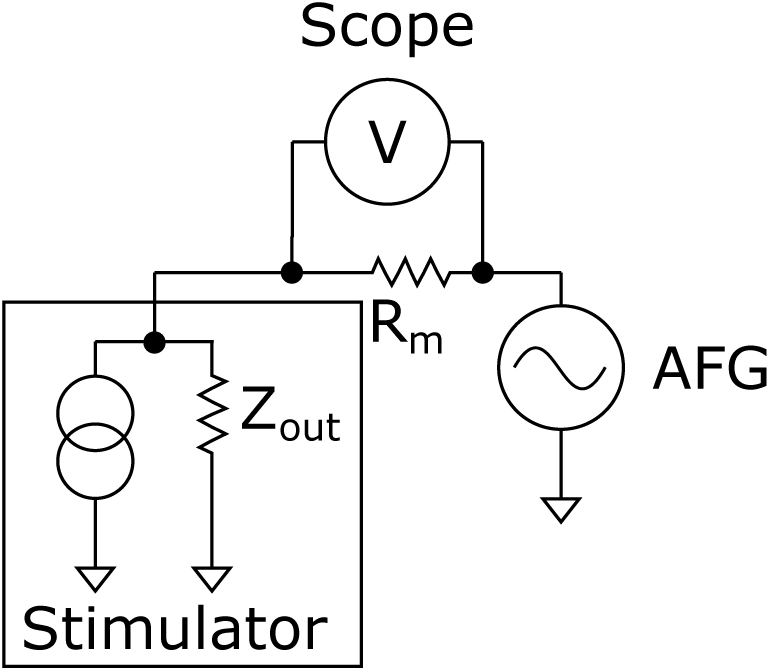
Instrument setup used to measure the output impedance of the Howland current pump.

### 3.2. Resolution and Linearity

To measure system resolution and linearity, all DAC output values were swept and the resulting output current measured through a 819 Ω resistor using a 434-series Wavesurfer Oscilloscope (LeCroy). A resistor below 1 kΩ had to be used in order to improve the measurement accuracy by using lower voltage per division and analog offset settings on the oscilloscope. Data was filtered by averaging at each step. Step size was on average −5.6784 µA/LSB. DNL and INL are plotted on Figure 8. As output displayed non-linearity at the extremes of the range the INL plot was cropped accordingly to reflect performance in the linear range of the circuit. The output range of the circuit spans *±*10 mA as shown in Figure 7.

**Figure 8:**
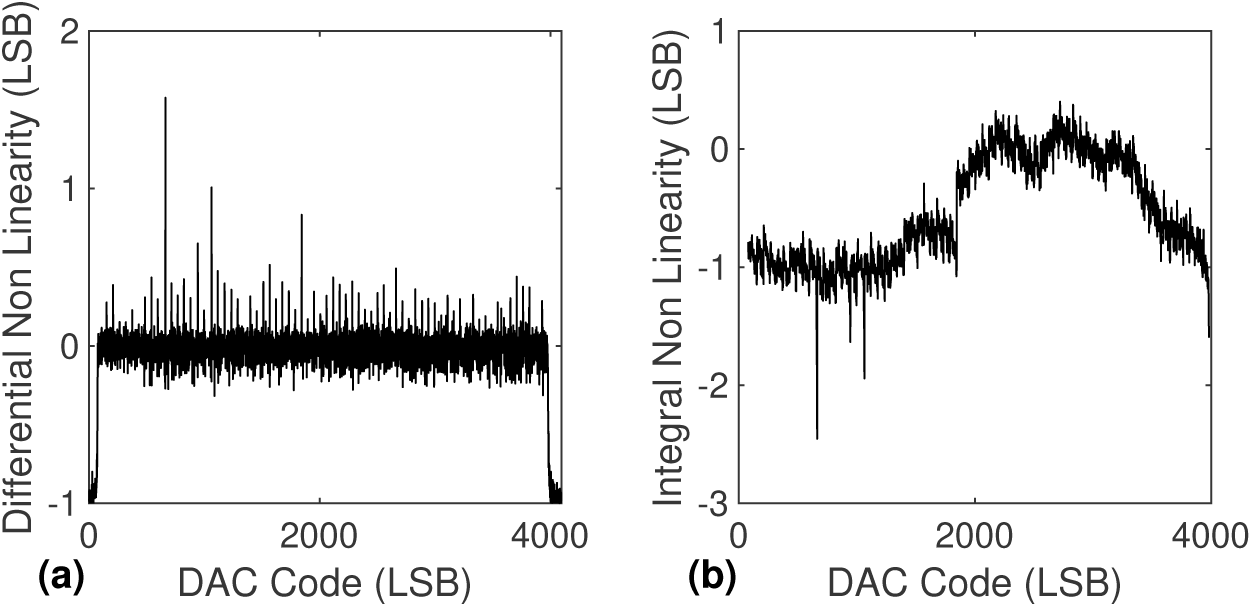
(a) Differential non linearity (DNL), normalised to LSBs; (b) Integral non-linearity, normalised to LSBs. Note that the INL plot in (b) has been cropped to display values in the linear operating range of the circuit.

## 4. Acute *Ex-Vivo* Experimentation Results

### 4.1. Ethics and Veterinary Surgery

All animal care and procedures were performed under appropriate licences issued by the UK Home office under the Animals (Scientific Procedures) Act (1986) and were approved by the Animal Welfare and Ethical Review Board of Imperial College University.

### 4.2. Protocol

Sprague-Dawley rats weighing 250-450 grams were used in this study. Briefly, animals were initially anaesthetised in a box with 5% isoflurane in oxygen, then culled by cervical dislocation prior to dissection of the sciatic nerve. The entire length of the nerve from the medial side of the ankle to the cleft close to the spine was used.

The sciatic nerve was placed in iced modified Krebs-Henseleit buffer and carefully cleaned of residual connective, muscular and vascular tissue. Superficial branches were cut close to the main trunk of the nerve such that only two branches extending the complete length of the sample remained. The final sample length is approximately 5 cm.

The nerve is then placed in a perfused dual chamber nerve bath. One partition is perfused with warmed and oxygenated buffer to preserve homeostasis and tissue viability. A 1 mm diameter hole allows threading of the distal end of the nerve into the second chamber which is filled with mineral oil, as shown Figure 9. Any leak between the two chambers is prevented using silicone grease applied using a large gauge syringe. A pair of silver-silver chloride electrodes spaced 2 mm apart are used in the oil partition for recording through a differential low noise amplifier (SR560, Stanford Research Systems). In the saline partition an 800 µm inner diameter conformal nerve cuff (Cortec) and an 800 µm inner diameter custom-made conductive elastomer nerve cuff[36] were implanted for stimulation and block of the nerve respectively. For current return a large 2 cm side square sheet of platinum is used and connected to stimulator ground.

**Figure 9:**
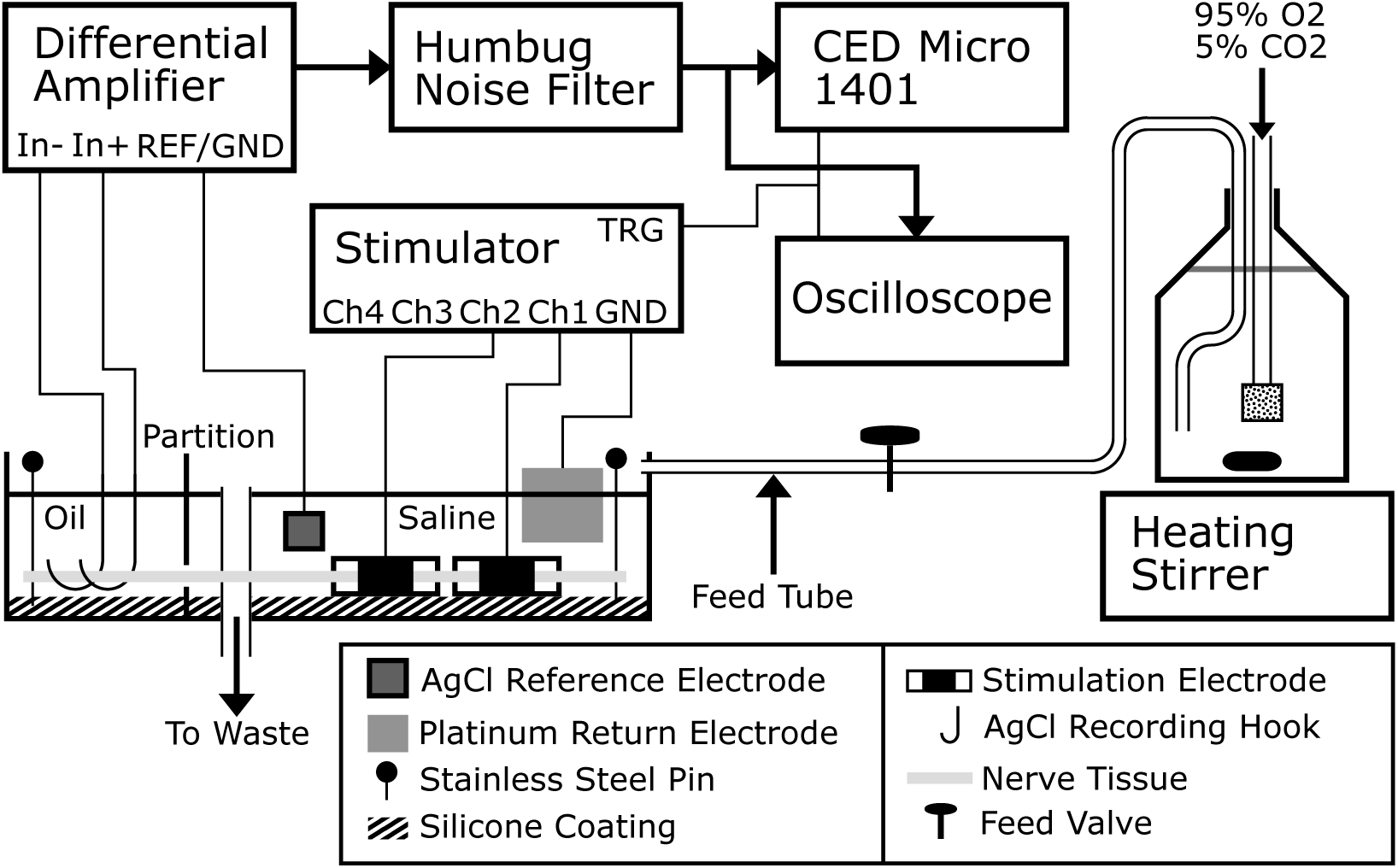
Drawing of the experimental setup for stimulating and recording CAPs from dissected nerve tissue *ex-vivo*. The CED Micro 1401 which digitises the analog signal and the stimulator itself are controlled by an external computer, not shown. The stimulator TRG signal is used to trigger both the oscilloscope and CED Micro 1401 acquisition hardware to timestamp stimulation events. A simple siphon is used to provide fresh buffer to the bath, while a peristaltic pump is used to drain the bath to avoid irregularity when using gravitational drainage. Two stimulation electrodes are used in tests where HFAC block is applied, and one when only conventional stimulation is required. The platinum sheet electrode is a common return path for current sourced through every channel of the stimulator.

### 4.3. Stimulation Capability

The ability of the stimulator to evoked compound action potentials (CAPs) in the A and C-type fibres of the rat sciatic nerve was evaluated using the aforementioned setup. In an example experiment, activation of A fibre and C fibres, or A fibres only, was possible by changing stimulation amplitude as shown Figure 10. Stimulation was carried out by using a cathodic-phase-first biphasic symmetric pulse of either 1000 µA or 3000 µA amplitude and 300 µs duration with a 20 µs interphase.

**Figure 10:**
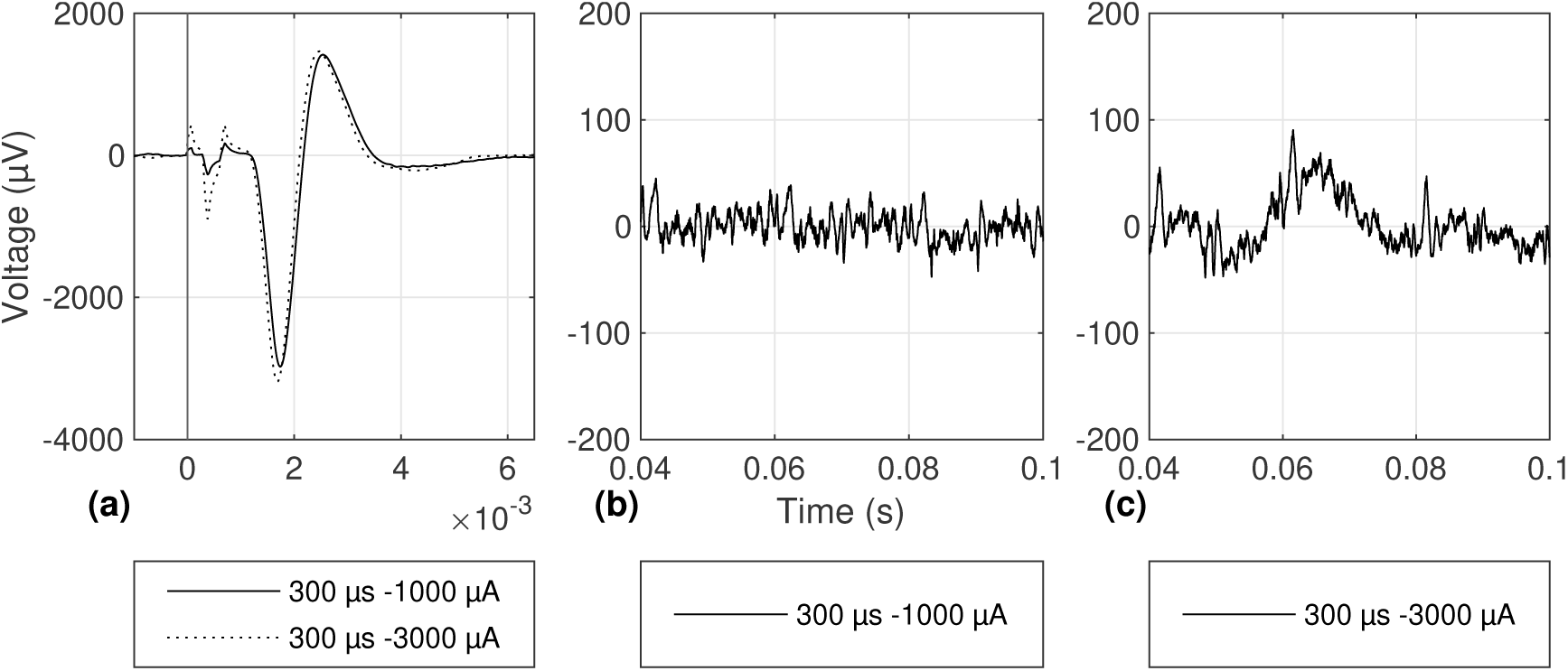
Recorded CAPs showing nerve response at different stimulation parameters and at different times after the stimulation trigger: (a) A fibre CAP for two different stimulation parameters, vertical line showing the start of stimulation; (b) neural waveform recording between 40 and 100 ms after the stimulation event for a biphasic symmetric pulse of 1000 µA amplitude and 300 µs duration; and (c) neural waveform recording between 40 and 100 ms after the stimulation event for a biphasic symmetric pulse of 3000 µA amplitude and 300 µs duration. Note the presence of a C fibre-type CAP.

### 4.4. Block Capability

Blocking performance for the stimulator was evaluated in the rat sciatic nerve model using the setup described in Figure 9. A and C fibre recruitment and nerve viability baseline was first measured by stimulating the sciatic nerve with 3*×* 2500 µA amplitude, cathodic first, biphasic symmetric, current controlled pulses. After three pulses were delivered HFAC block was applied at the stimulation electrode between the stimulating and recording electrodes, corresponding to stimulator channel 2 on Figure 9. Block was applied as a current-controlled square wave at 10 kHz and 6 mA amplitude while stimulation continued using the same parameters at the rate of 1 Hz during block, as shown on Figure 11. Block was then terminated after 15 seconds and stimulation continued for 7 additional pulses to evaluate nerve recovery from block. C fibres appear to recover more slowly from block than A fibres as can be seen during the recovery phase, despite the fact that they are only partially blocked at this block amplitude and frequency in this trial. It was not possible to obtain complete block of C fibres during the stimulator test.

**Figure 11:**
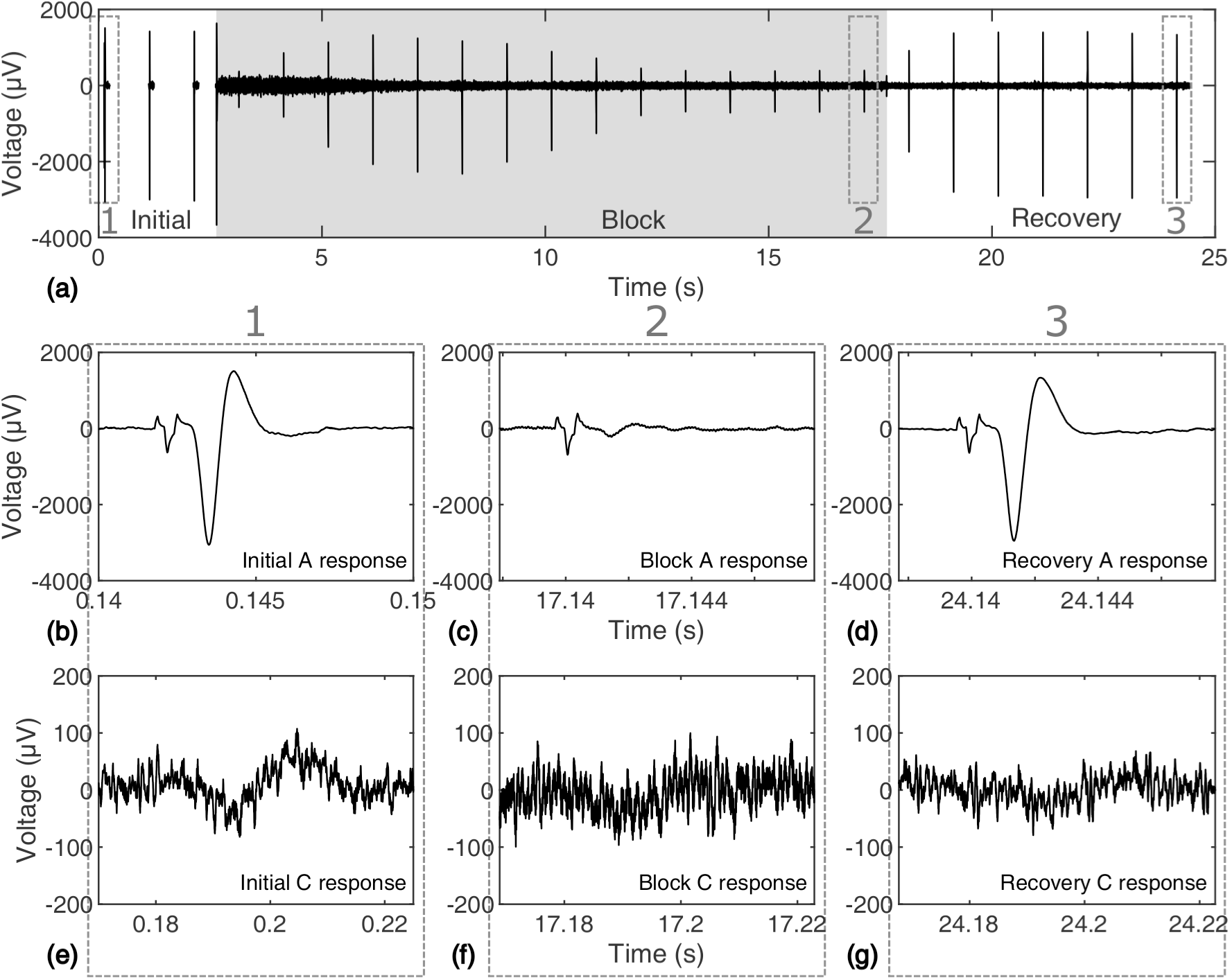
Block and recovery trial in which A fibres are completely blocked and C fibres are partially blocked: (a) overview of the trial and timeline starting with initial stimulation without block, application of block and continuing stimulation, and cessation of block with nerve recovery; (b) initial A fibre response; (c) A fibre response at the end of the blocking phase; (d) A fibre response at the end of the recovery phase. (e) initial C fibre response; (f) C fibre response at the end of the block phase; and (g) C fibre response at the end of the recovery phase.

## 5. Discussion

Achieved specifications for the stimulator with respect to other designs are shown Table 3, however on their own they do not completely describe its strengths and weaknesses.

**Table 3:** Measured performance specifications for the stimulator compared to other similar designs.

| Parameter | [1] | [28] | [2] | [37] | this work |
| --- | --- | --- | --- | --- | --- |
| Device Type | CMOS | CMOS | COTS | COTS | COTS |
| Compliance Voltage (V) | N/A | 17 | 18 | 120 | $\pm 15$ |
| Full-scale Current (mA) | 1 | 0.31 | 0.250 | 10 | $\pm 10$ |
| Current Resolution (μA) | N/A | 10 | 1 | analog | 5.6784 |
| Stimulation Timing Resolution (μs) | N/A | 0.8 | 2 | 5 | 4 |
| Channel Number | 4 | 8 | 1 | 1 | 4 |
| Power Consumption (W) | 0.0189 mW | 0.029 | 0.165 | 1.8 | $\leq 4$ |
| Size (cm) | $3 \times 1.5 \times 0.5$ | N/A | $2 \times 4 \times 0.5$ | N/A | $9 \times 6 \times 5$ |

The key strengths of the proposed stimulator design are:

(i) Based on Commercial Off-The-Shelf (COTS) components.
(ii) Cheap to source and assemble (less than GBP 350 or USD 450).
(iii) Flexible powering options, including the controlling computer through a dedicated 5W USB port.
(iv) Complete control of output waveform, scripted using any program that can interface with the FTDI Virtual COM Port driver.
(v) High current output combined with high resolution and voltage compliance.
(vi) Multiple independent channels that are driven with accurate timing.
(vii) Feature extendability using additional PCB modules, such as for recording, that can be driven by the microcontroller processor.

While the device allows concurrent current-controlled multi-channel stimulation, this is only possible when stimulation is delivered in a monopolar configuration with a common return electrode connected to stimulator ground, which must be placed away from the nerve in the bath to prevent interference during stimulation. It is possible to have completely isolated current output channels on one device, however this requires individual isolated high-voltage supplies for each channel with their own battery, 12-bit DAC and digital isolator. The added complexity was deemed too high due to the added bulk, cost and power consumption. When several bipolar channels are absolutely required, several stimulators can be used in tandem and controlled using the same computer, each requiring a USB port and external battery.

The cutoff frequency for output impedance which is shown Figure 7 is low at only 1 kHz and can be attributed to the wide output swing required for U1 as a result of U2 being in a non-inverting configuration. However for the purposes of stimulation using cuff electrodes the output impedance is still much higher than the kilo-ohm range at all frequencies of interest, for example the impedance magnitude at 10 kHz is near 40 kΩ, leading to less than 3% error in current output for electrodes having 1 kΩ impedance at that frequency. It is however possible to shift the cutoff frequency to the right by adjusting the design, trading power by using an inverting configuration for U2 for example. The simulated and measured curves show general agreement yet differences suggest that careful optimisation could improve the AC performance of the stimulator incrementally, potentially gaining an order of magnitude of output impedance at the highest frequencies. Output compliance could be nudged upwards by the use of a rail-to-rail amplifier for U2, however there could be possible trade-offs in output impedance at high frequencies since U2 was chosen specifically for its high current output and high slew rate.

Currently power consumption is quite high at approximately 4 W measured using a DC Power Analyzer (N6705B, Agilent Technologies) compared to similar stimulators in the literature, though optimization of power was not a priority in this design due to the inherent power demand of high frequency block. Currently, using 5V USB-chargeable battery packs for power ensures both long operating time on one charge and straightforward access to charging. However there is room to optimize for power consumption both on the stimulator module itself by providing ways for the microcontroller to turn off stimulator channels that aren’t connected to electrodes, and also by optimizing the power consumption of the microcontroller itself. The former is possible by daisy-chaining additional switches to the SPI channel used for channel output routing, and the latter could benefit from a slower core clock, however this would affect timing performance. Idle power could be further reduced by gating the clock to select peripherals only when they are needed.

In terms of performance during electrophysiology experiments, both A and C fibres were able to be stimulated in the *ex-vivo* rat sciatic nerve model. While A fibre block was achieved C fibre block was only partial as can be seen Figure 10. There are several possible reasons for this including nonidealities in the design of the nerve cuffs used as well as possibly insufficient voltage compliance or insufficient current output to reach the high block thresholds of C fibres, which are higher than those of A fibres. Work is ongoing to identify the root causes of this observation. The stimulator also does not have a DC-filter to prevent DC leakage, which may have affected the block thresholds of C fibres. Leakage can be calibrated to be less than 5 µA and was verified to remain constant over the course of several hours. In future experiments to determine the block threshold of C fibres, an external DC filter can be used such as the one described in [24].

## 6. Ongoing and Future Work

Although the hardware is present for the stimulator to monitor electrode voltage during stimulation, software is still currently being developed. Work is ongoing to develop software to detect over-compliance during stimulation and measure the impedance of electrodes for characterization and diagnostics, for example.

On the PC side the current programming model is script-based using MATLAB, and work is ongoing to develop a GUI which will streamline the use of the device for concurrent stimulation and recording, and allow marking of stimulation events along with corresponding stimulation settings for each event, for example.

## 7. Conclusions

A stimulator design is proposed that aims to fulfill requirements for a versatile and powerful acute *ex-vivo* and *in-vivo* neurophysiology platform. The stimulator is suitable for both conventional and blocking stimulation of peripheral nerves, and was designed with versatility and affordability in mind. It was shown to be able to excite A and C fibre populations which represent both ends of the nerve fibre diameter spectrum in the rat sciatic nerve, and can be assumed suitable for other peripheral nerves such as the vagus. Complete control of the output waveform is given to the user combined with accurate timing using address-event representation for stimulation. Stimulation can be carried out using multiple channels concurrently for monopolar electrodes. Power is provided by USB-chargeable batteries to prevent interference of stimulation with recording.

The overall goal is to provide an affordable tool for neurophysiology experiments with a quick turnaround time to de-risk more involved chronic *in-vivo* studies, in light of interest in using HFAC block combined with conventional stimulation for therapeutic applications.

## Acknowledgements

The authors gratefully acknowledge Galvani Bioelectronics (Stevenage, United Kingdom) for providing an experimental opportunity to test a stimulator prototype, Dr Rylie Green and Dr Omaer Syed (Imperial College London, Department of Bioengineering) for assistance with animal experiments, and Dr Lieuwe Leene (Imperial College London, Department of Electrical and Electronic Engineering) for help with debugging the Howland Current Pump design. Special thanks to Dr G.E Hunsberger (Galvani Bioelectronics), Dr W. Grill and Dr N. Pelot (Duke University Biomedical Engineering) for help with troubleshooting the *ex-vivo* neurophysiology setup used to test the proposed stimulator.

AR is supported through the EPSRC Centre for Doctoral Training (CDT) in High Performance Embedded & Distributed Systems (HiPEDS, EP/L016796/1).

## Open Access Statement

All design resources (including PCB design files, firmware, Matlab code) will be made freely available, either through our institutional website www.imperial.ac.uk/next-generation-neural-interfaces/resources, github, or by directly contacting the authors.

